# Pan-African model explains *Homo sapiens* genetic and morphological evolution

**DOI:** 10.1101/2025.05.22.655514

**Authors:** Cecilia Padilla-Iglesias, Zhe Xue, Michela Leonardi, Manolo Fernandez Perez, Johanna L.A. Paijmans, Margherita Colucci, Anahit Hovhannisyan, Pierpaolo Maisano-Delser, Javier Blanco-Portillo, Alexander G. Ioannidis, Giulio Lucarini, Jacopo N. Cerasoni, Andrew W. Kandel, Manuel Will, Emily Y. Hallett, Mario Krapp, Karen Lupo, Eleanor M.L. Scerri, Isabelle Crevecoeur, Lucio Vinicius, Andrea B. Migliano, Andrea Manica

**Affiliations:** Evolutionary Ecology Group, Department of Zoology, University of Cambridge, Downing Street, Cambridge, CB2 3EJ, UK; Emmanuel College, University of Cambridge, St Andrew’s St, Cambridge, CB2 3AP, UK; McDonald Institute for Archaeological Research, University of Cambridge, Downing St, Cambridge CB2 3ER, UK; Human Evolutionary Ecology Group, Department of Anthropology, University of Zurich, Winterthurerstrasse 190, 8057, Zurich, Switzerland; Natural History Museum, Cromwell Road, London SW7 5BD, United Kingdom; Department of Biodiversity and Conservation, Real Jardín Botánico (RJB-CSIC), Madrid, 28014, Spain; School of Environmental and Natural Sciences, Bangor University, Deiniol Rd, Bangor LL57 2UR, UK; Human Palaeosystems Research Group, Max Planck Institute of Geoanthropology, Kahlaische Strasse 10, Jena, 07745, Germany; Smurfit Institute of Genetics, Trinity College Dublin, Dublin D02 PN40, Ireland; Department of Biology, Stanford University, 371 Jane Stanford Way, Stanford, CA 94305, USA; Department of Biomolecular Engineering and Genomics Institute, University of California, 1156 High Street, Santa Cruz, CA 95064, USA; Department of Biomedical Data Science, Stanford Medical School, 300 Pasteur, Edwards, Palo Alto, CA 94304, USA; National Research Council of Italy, Institute of Heritage Science (CNR-ISPC), Piazza Guglielmo Marconi, 10, Rome 00144, Italy; Department of Biology, Loyola University Chicago, 1032 W. Sheridan Road,Chicago, IL 60660, USA; The Role of Culture in Early Expansions of Humans (ROCEEH), Heidelberg Academy of Sciences and Humanities at Eberhard Karls University of Tübingen, Hölderlinerstrasse 112, 72074 Tübingen, Germany; Department of Early Prehistory and Quaternary Ecology, University of Tübingen, Burgsteige 11, 72070 Tübingen, Germany; Palaeo-Research Institute, University of Johannesburg, P.O. Box 524, Auckland Park, ZA-2006, South Africa; Department of Anthropology, Loyola University Chicago, 1032 W. Sheridan Road, Chicago, IL 60660, USA; GNS Science, 82 Wyndham Street, Auckland CBD, Auckland 1010, New Zealand; Department of Anthropology, Southern Methodist University, PO Box 750336, Dallas, TX 75275, USA; Department of Classics and Archaeology, University of Malta, Msida MSD 2080, Msida, Malta; Department of Prehistory, University of Cologne, Kerpener Strasse 30, 50937Cologne, Germany; University of Bordeaux, CNRS, Ministère de la Culture, PACEA, UMR 5199, F-33600 Pessac, France

## Abstract

A growing body of evidence has challenged the traditional assumption of a single-region origin for *Homo sapiens*, suggesting instead that our species originated from multiple geographically distinct populations in Africa, which intermittently exchanged genes and culture. However, our understanding of how this Pan-African metapopulation would have evolved through time is still limited. Furthermore, the drivers of such changes are uncertain, and quantitative models of the respective contributions of different African regions are lacking. Here we provide a complete reconstruction of the meta-population dynamics over the last 200,000 years by quantitatively integrating an ecological niche model, informed by archaeological sites, within a spatially explicit population genetic framework. The inferred metapopulation dynamics account for the divergence among all available contemporary and ancient genomes of African hunter-gatherers used to calibrate the model. In addition, it also accurately predicts the patterns of craniometric diversification across the continent from the Late Middle Pleistocene to the present. Finally, we show how the climate-driven changes in population sizes and connectivity are congruent with major patterns of archaeological and phenotypic diversification over the last 200,000 years across the African continent.

## Main text

Over the past few decades, a large number of studies have revealed an African origin for all non-African populations, dating between 70 and 50 thousand years ago (ka)^1–5^, and much progress has been made in documenting the factors that might have promoted or facilitated such a dispersal^3,6–8^. Comparatively less is known about the origins, geography and diversification of *Homo sapiens* in Africa. While a single African population origin was postulated until recently, current research in paleoanthropology, paleoecology, archaeology and population genetics has suggested that humans likely evolved within a subdivided African metapopulation, maintained by genetic and cultural exchanges influenced, at least in part, by environmental conditions^2,9–12^.

Genetic studies have been instrumental in confirming the contribution of multiple African populations to the biological diversity of modern humans^13–19^. Models of genetic evolution among contemporary and ancient hunter-gatherers in Africa suggest a mix from 3-4 stem populations that diverged from one another between 300-100 ka^2,13,14,17,18,20–27^. Yet, our knowledge of the geographical and demographic patterns characterising these lineages during the early evolution of *H. sapiens* is extremely limited. Today’s African hunter-gatherers show a closer genetic affinity to ancient genomes obtained from regions in and around their geographic range than to other hunter-gatherer populations elsewhere in Africa. However, no ancient genomes in Africa are older than 15 ka^14,16,23,28,29^, and there are no extant hunter-gatherers in Northern and Western Africa. Evidence of population structure in the deep past mostly stems from studies looking at archaeological and fossil data. First, the defining characteristics of *H. sapiens* do not appear in the fossil record at a single location or time in Africa, but rather as a mosaic-like mix of primitive and derived features in different African regions^9,10,30,31^. Similarly, the Middle Stone Age (MSA, ranging from about 300-30 ka; although it differs by region), the first and longest lasting cultural stage associated with *H. sapiens* appears simultaneously in more than one African region, rather than radiating outwards from a single source in the Late Middle Pleistocene^9,32^. Still, many questions regarding the origins, geographical boundaries, and dependence on environmental factors of a hypothesised long-standing African metapopulation remain unanswered^18,26,33–35^.

Here we provide the first comprehensive reconstruction of the Pan-African metapopulation dynamics over the last 200,000 years by quantitatively integrating genomic data (Fig 1A; Dataset S1), the location and dates of archaeological sites (Fig S1; Dataset S2), and palaeoclimate simulations within a Climate Informed Spatial Genetic Modelling (CISGeM) framework^8,36^ (Fig S2). We started by applying a time-dependent Ecological Niche Model (ENM^37,38^) to quantify the bioclimatic niche of *H. sapiens* over time based on the distribution of N=1242 dated archaeological assemblages on the continent which attest to definite human occupation (Fig S1-S3; Dataset S2). A time-dependent ENM provides a diachronic estimation of the climatic conditions associated with observations and allows the reconstruction of the changes in geographic range of suitable areas through time. We then used the resulting ENM suitability scores through time and space as inputs for CISGeM (Fig S2). CISGeM simulates environmentally-driven changes in population size, mobility and range, and matches them to observed genetic diversity, in this instance among contemporary and ancient African hunter-gatherers. We used a Monte-Carlo approach to sample basic demographic parameters, such as the link between ENM suitability scores and local population sizes, population growth rate and migration, and the impact of altitudinal changes on migration. We then fitted the parameters using a Deep Learning (DL) approach based on a Convolutional Neural Network (CNN)^39^ architecture using five summaries (mean, median, maximum, standard deviation and median absolute deviation) calculated from the matrices of pairwise genetic differentiation (**π**) between all possible pairs of populations over 2,000 genomic windows (see Materials and Methods).

**Figure 1.**
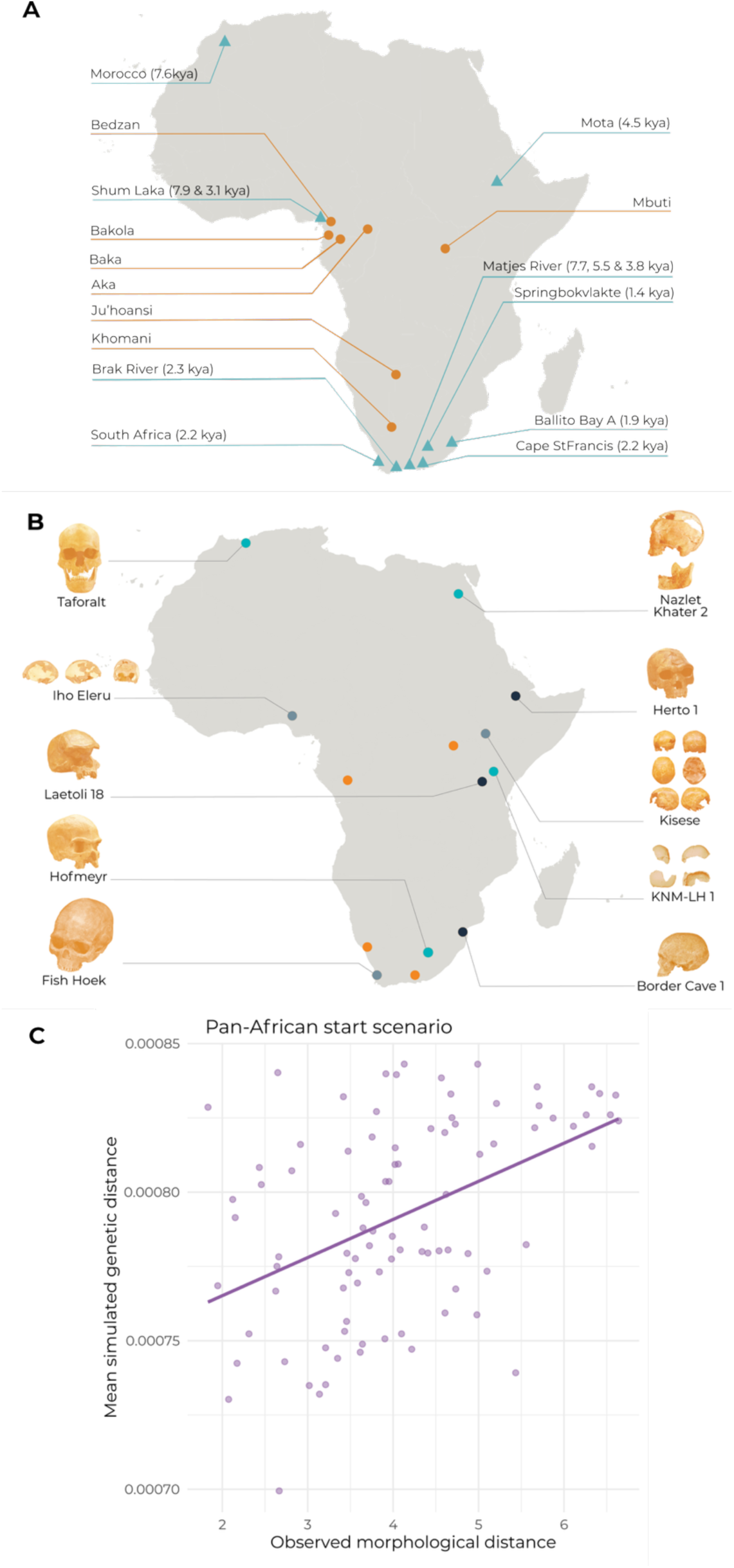
Genetic and morphological samples included in our study. A) Map of samples used for running CISGeM, where orange dots indicate contemporary populations and blue triangles indicate ancient genomes. Names denote the name given to the genome, rather than the archaeological site. B) Cranial specimens (grouped by population) included in our analyses, with orange dots indicating contemporary samples, grey dots corresponding to Holocene samples, light blue denoting Late Pleistocene samples, and black denoting Middle Pleistocene samples. C) Relationship between observed morphological distances between fossil populations and mean simulated genetic distances at their respective locations and time periods for the Pan-African start scenario. The blue line represents a linear model least squares regression, and dots correspond to each pairwise comparison between fossil populations.

### A continuous Pan-African metapopulation since 200 ka reconstructs African genetic diversity

Fossils that arguably display close morphological affinities to *H. sapiens* first appear during MIS (Marine Isotope Stage) 8 (300-243 ka)^40^, with notable sites including Florisbad in South Africa^9,41–45^, Jebel Irhoud in Morocco^31,40,46^, and the Kibish Formation in Ethiopia^9,47,48^. These fossils are also associated with early examples of the MSA. For this reason, we fitted four scenarios in CISGeM with a different geographical origin: the Kibish Formation (henceforth Eastern African start), the Jebel Irhoud site (Northern African start), the Florisbad site (Southern African start), and finally, a model assuming a population already distributed across the entire continent according to values from our ENM (Pan-African start)(Table S1).

The simulated scenarios were able to recreate the patterns of genetic differentiation across the continent. The range of observed values for all comparisons (161) of the five pairwise ***π***-based summary statistics were within the simulated ranges (Fig S5-S6). Even when we focus on each of the 161 between-population average pairwise observed ***π*** values individually, all of them fall within the simulated range (Fig. S5). Moreover, we observed a high classification performance for model selection, with no individual class accuracy below 87% (Fig S7). When we compared the relative support for each of the four scenarios using DL, we found that the Pan-African start scenario showed the highest support (0.519 probability), followed by the Southern and Northern African start scenarios with comparable support (0.263 and 0.205 probability, respectively), and by the Eastern African scenario with much lower support (0.012). After confirming that our model had sufficient power to estimate parameter values (Table S2), we used DL to generate posterior probabilities of the input demographic parameters given the observed levels of genetic differentiation. Our inferred initial time for the simulation (representing the point from which metapopulation structure is introduced in the model) tended to cluster around 11,000-9,000 generations ago (275,000-225,000 years) in the Pan-African, Northern African and Southern African start scenarios, whilst it did so around 12,000-11,000 generations ago (around 300,000-275,000 years) in the Eastern African scenario (Fig S8-S11; Table S3). This timeframe coincides with estimates of the earliest divergence of human genetic lineages. The divergence of San hunter-gatherers of southern Africa from all other human genetic lineages is thought to have occurred around 270 ka (350-260 ka)^21^ and of Central African hunter-gatherers from all other human genetic lineages at ca. 221 ka^17^, but see^14,34,49^. It also coincides with the appearance of the first fossil remains showing a mosaic of anatomically modern and archaic characteristics including Irhoud 1, Jebel Irhoud, Morocco (315 ka)^31^; KNM-ES 11693, Eliye Springs, Kenya (285±15 ka)^50^; LH 18, Laetoli, Tanzania (250±50 ka)^51,52^, Florisbad (259±35 ka)^43^, Omo Kibish 1 and 2, Ethiopia (233± 22 ka)^9,46–48^ and the emergence of MSA technologies in these regions^10,31,53,54^. Across scenarios, we also obtain an effective size for the ancestral population within the range of 15,000-18,000 individuals, which matches other genetic estimates^34^.

Next, we selected the 1000 best fitting runs of all the scenarios based on the sum of Euclidean distances between observed and simulated summary statistics to reconstruct the demography and migration patterns of *H. sapiens* through space and time (Fig S12-S20). Despite their distinct geographical starting points, from 200 ka the simulated demographies were very similar under the four start scenarios (Table S4; Pearson *r*>0.96-0.99, *p*<0.001 between all pairs of scenarios), indicating that, after an initial, scenario-specific phase setting up the appropriate genetic diversity across the continent, the model converged to a relatively consistent metapopulation dynamics up to the present. We then used these reconstructions to validate the model against anthropological data not used when fitting the model. We did so by first comparing our predicted population densities for the present across the continent with an equivalent map of densities based on ethnographic censuses from N=358 hunter-gatherer societies around the world (N=21 African)^55^, and found a high correlation (Spearman rho=0.53, *p*<0.001; for all scenarios irrespective of the starting location)(Fig S21). Second, using a compilation of N=1,311 camp locations from contemporary hunter-gatherer societies in Central, Eastern and Southern Africa^56^, we verified that predicted effective population densities at hunter-gatherer camps were significantly higher than at randomly sampled locations (Wilcoxon rank-sum tests, *p*<10^-25^, irrespective of the start scenario), with a medium effect size (Cliff’s δ=0.38), indicating that the model successfully identifies areas suitable for hunter-gatherer occupation (Fig S22).

### Simulated demographic histories match H. sapiens morphological differentiation

We further validated our metapopulation model by demonstrating that it can predict morphological differentiation from 200 ka to the present across the continent, using 45 crania from 14 populations (Dataset S3; Fig 1B-1C; Fig S23). We used the same best-fitting 1000 simulations from each of the start scenarios to estimate the expected genetic differentiation of populations inhabiting the geographical location of the crania at their respective dates, and compared such expected genetic differentiation to pairwise morphological distance based on 13 size-adjusted, putatively neutral cranial measurements^57,58^. We obtained a strong correlation between the mean simulated genetic distance and the observed morphological distance (Mantel tests for the Pan-African, Southern, Northern and Eastern start: r=0.464, *p*=0.002; r=0.461, *p*=0.016; r=0.391, *p*=0.008; r=0.454, *p*=0.008; respectively). The strength of this relationship is in line with the expected relationship between cranial phenotypes and genetics in humans^57,58^. Moreover, it could not be explained merely by isolation by distance. We tested this by verifying that observed morphological distance was not correlated between two different measures of geographic distance: simple great-circle distances and distances cost-adjusted for topography (Table S5) and by checking that controlling for such measures of distance in partial Mantel tests did not alter the correlation between simulated genetic distance and morphological distance (Table S6). Therefore, our simulated climate-driven changes in population sizes and migration over time, which were fitted on genomes from contemporary and relatively recent (<10 ka) hunter-gatherers, also predict the broad-scale patterns of morphological diversification in *H. sapiens* across Africa from 200 ka.

### Pan-African demographic histories identify key regions for H. sapiens evolution

Irrespective of the starting location of the simulations, the majority of Southern Africa, Eastern Africa and the Maghreb would have remained populated from MIS 7 (at around 200 ka) until the present. These regions have been repeatedly highlighted as the key areas where derived innovations characterising *H. sapiens* first occurred^59–61^, and it is notable that our model recovers their long-term demographic persistence independently of those early fossil and archaeological observations, as the ENMs underlying CISGeM were trained only on archaeological sites up to ∼120 ka. Whilst the archaeological record of Central and Western Africa is less complete and significantly more disturbed (and probably not representative of the full behavioural range of these populations^62^), our model suggests that Southern Africa, Eastern Africa and the Maghreb might have been exceptional in their ability to maintain sufficiently large or dense population sizes over long periods for adaptive innovations to repeatedly emerge, and to remain sufficiently connected to facilitate their spread (Fig. 2).

**Figure 2.**
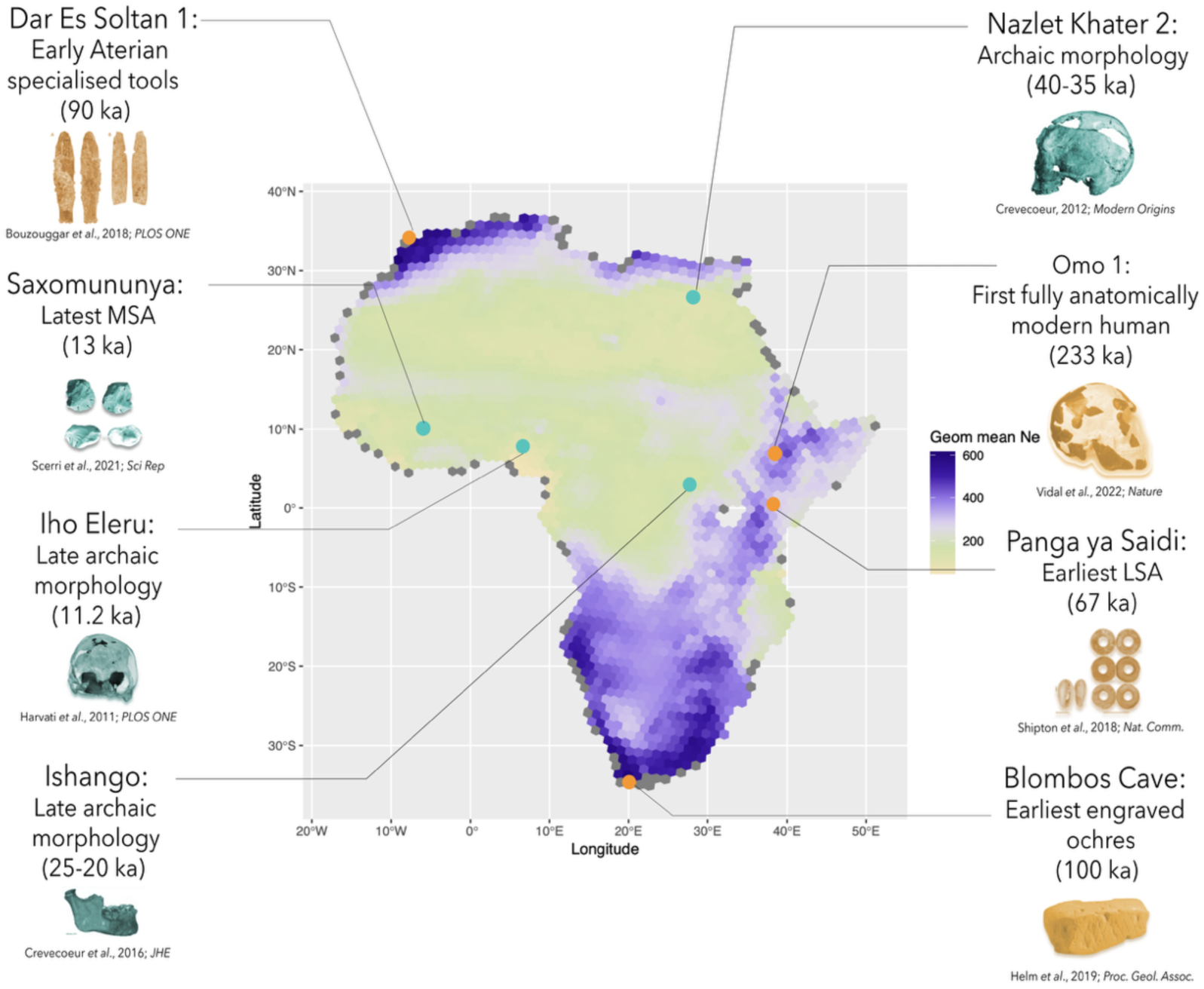
Variability in demography throughout *Homo sapiens* evolutionary history. Geometric mean of effective population size per CISGeM deme across all time periods across the 1000 best fitting runs of the Pan-African start scenario. Grey dots represent demes that remain uninhabitable for over 50% of time-steps due to changes in sea levels. Blue dots represent examples of archaic morphological or cultural traits, whilst orange ones depict examples of morphological or cultural innovations.

Similarly, the reconstructed repeated gains and losses of connectivity across the Sahara match the peculiarities of both the morphological and archaeological record of Northern Africa dating between 150 ka and 30 ka^30,63^. Despite the early appearance of archaic *H. sapiens* in the region^46^, later fossils such as El Harhoura 1 (66±5 ka), Contrebandiers I (103±8 ka), or Dar es Soltan II (80±5 ka)^30,63^ show characteristics that do not resemble other sub-Saharan specimens from similar time periods. However, these fossils show a similar retention of archaic morphological traits, which together with the widespread and enduring presence of Aterian industries across the Maghreb, suggest a degree of regional continuity and connectivity^30,46,64^. On the contrary, our model reveals much higher levels of demographic instability and connectivity losses in Egypt, both with the Maghreb and sub-Saharan Africa, at multiple times. In that region we observe recent fossils that still retain archaic features, such as the remains found at Nazlet Khater 2 (NK2, 38±6 ka^65^), or more recent Holocene specimens that also show robust phenotypes hypothesised to be related to NK2^66^, and result from the Nile valley’s role as a refuge area^67,68^.

Our model reveals that West Africa, and in particular the area around the Gulf of Guinea as well as Senegal, is the region with the highest reconstructed population turnover on the continent and remains isolated for large periods throughout our evolutionary history until the recent past. This is consistent with fossil and archaeological data. First, the oldest yet known fossils in West Africa, while dating to as late as 11.2 ka (or 13 ka calibrated) in Iho Eleru, Nigeria^69,70^, show very archaic features (and outside of contemporary human variation). Second, this population is associated with some of the first Later Stone Age (LSA) industries in West Africa, which coexists with a late persistence of the MSA in this region—well after the MSA had been replaced by the LSA in Southern Africa, Eastern Africa and the Maghreb^71^. Genetic studies have shown that Southern African Stone Age hunter-gatherers (including those from our sample) despite representing the most diverged human genetic lineages, still share many more alleles with eastern Africans (including the present-day Dinka and Mota) than with present-day Western Africans (as represented by the Yoruba)^17,23,29^. Alongside the presence of some “archaic” morphological features and continued MSA industries in Western Africa in the Holocene, these findings suggest the existence of archaic lineages with no direct descendants in the region^17,21,23,69,72^. However, an alternative explanation is that the population cline encompassing Eastern and Southern Africa was not strongly connected to Western Africa. Accordingly, our demographic reconstructions show reduced levels of East-West migration compared with East-South migration for the vast majority of our evolutionary history, similar to what is observed in other mammals^73^. Our close prediction of the morphological distance between the Iho Eleru specimen and the rest of the cranial specimens in our sample provides additional evidence for our proposed scenario, though we do not exclude the possibility of small amounts of introgression.

We designed an additional test to determine whether a deep history of population interconnectivity could explain the genetic diversity observed in Western Africans. Although there are no extant hunter-gatherers in the region, the Yoruba represent a relatively “unadmixed” Western African population not affected by a large-scale farming expansion^17^ (Fig S24) and thus expected to bear strong genetic similarities to their hunter-gatherer ancestors. We took a sample of 20 high-coverage Yoruba genomes and used our 1000 best-fitting simulations to estimate where a population with their genetic profile would be currently placed by the model (Fig S25). The model correctly located the Yoruba within a narrow area of Western-Central Africa corresponding to their current range (Fig S26), indicating that a long history of environmentally-driven changes in population dynamics can explain Western Africa’s genetic, cultural and morphological evolutionary trajectories.

### Changes in population density and connectivity match key events in modern human evolution

After quantitatively validating our Pan-African metapopulation model of *H. sapiens* evolution with independent archaeological, anthropological, morphological and genetic evidence, we used the reconstructed population sizes and connectivity across the continent to contextualise key human evolutionary events from published literature over the last 200 ka (see Table S7 for a list of main events). We stress that congruence between large-scale demographic patterns and observed archaeological and phenotypic patterns does not imply causation but rather helps us interpret the possible contribution made by the simulated biological processes (demographic and genetic). Furthermore, the African archaeological record suffers from uneven sampling and variable resolution, meaning that absences of certain features should be interpreted with caution. Despite these caveats, we find that our demographic reconstructions are coherent with most of the events in Table S7, thus providing useful context for their interpretation. There are also several exceptions (Table S7); this is not surprising, since we expect other processes beyond demography to affect cultural transmission and phenotypic evolution. Thus, these exceptions should not be seen as invalidating the quantitative model, but rather provide fruitful direction for future work. Below we discuss the major patterns in chronological order but refer the reader to Table S7 for a more comprehensive assessment of the congruences and exceptions through space and time.

From MIS 7 onwards, a high level of morphological diversity is observed among fossil hominins within and between African regions^48,74,75^. Our model’s predicted deep population structure provides a potential mechanistic explanation for such diversity (see Fig. 3 for effective population sizes across African regions, and Fig. 4 for the reduced levels of between-region migration, shown in orange). Our model also provides support for the establishment of a stable corridor connecting Eastern and Southern Africa by 200 ka that remains uninterrupted for most of the period (see prevalence of high migration, blue colours, in the region in Fig. 4; and the large geometric mean of *Ne* across time in Fig. 2), as hypothesised by Skoglund et al.^23^; and Lipson et al.^14^ and as observed among other African mammals^76^. This corridor, alongside the extremely high levels of intra-regional connectivity predicted by our model in Southern Africa from MIS 6 onwards (blue colours in Southern Africa in Fig. 4), also matches the early manifestations of anatomically modern human morphology in Southern Africa, e.g. several remains in Klasies River (100±10 ka), Border Cave BC5 (74±5 ka) and BC1 (126±44 ka) specimens^77–79^. Moreover, although our model does not make quantitative predictions about cultural variation or transitions, it offers relevant insights into their demographic contexts. For instance, high levels of connectivity in Southern Africa for most of the study period can account for the early and continued appearance in the Late Pleistocene of complex, cumulative cultural traits and behavioural innovations in the region. These are features typical of later *H. sapien*s and incorporate the widespread use of pigments including abstract engravings, systematic exploitation of marine resources, and the production of shell beads and bone tools^80,81^.

**Figure 3.**
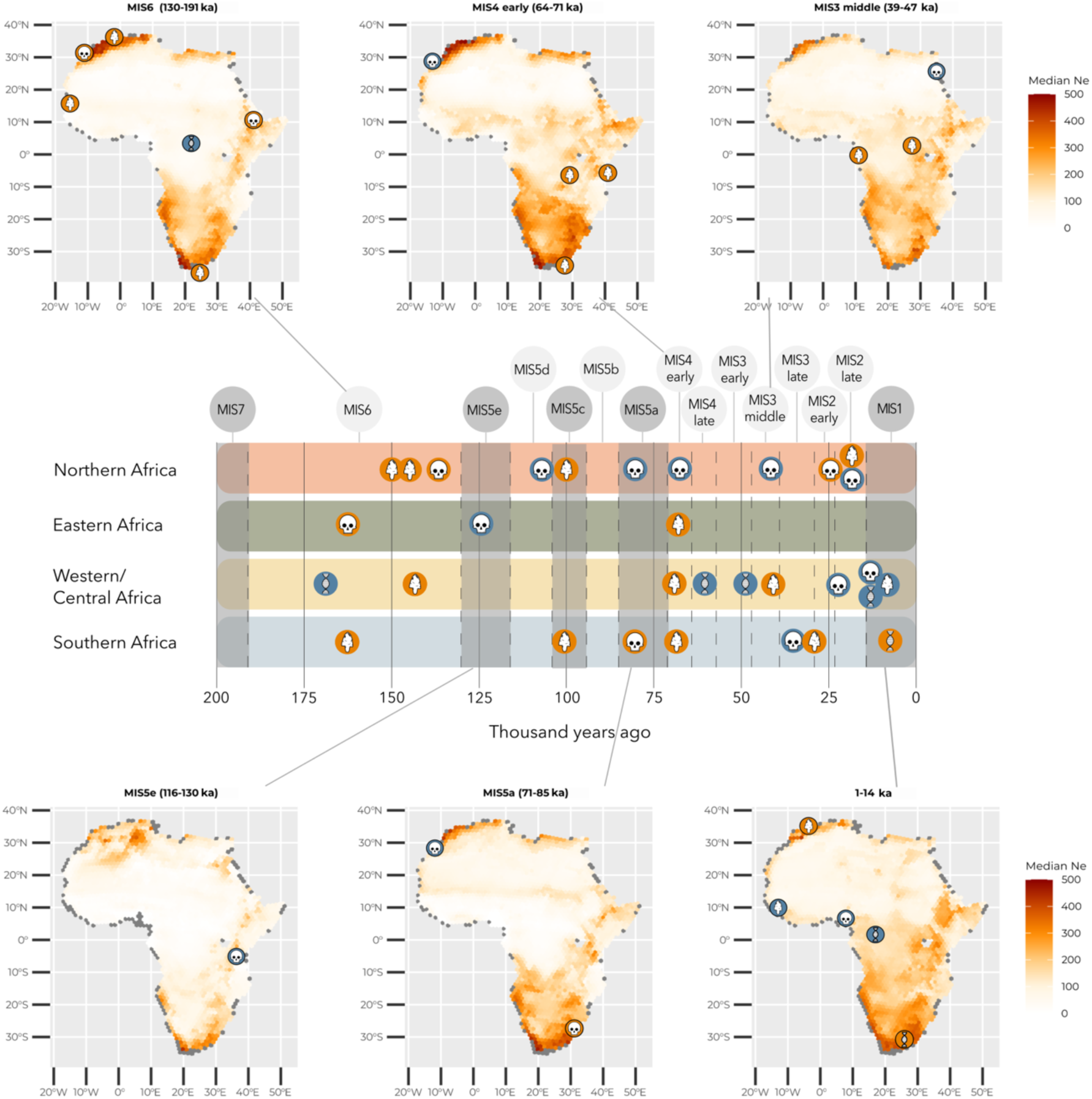
Demography over time and key events within regions. Maps are median effective population sizes for each cell at each MIS stage for the Pan-African start scenario. Timeline indicates key genetic (DNA symbol), archaeological (stone tool symbol) and morphological (skull symbol) events occurring in different African regions (see Table S7). Interglacial periods shaded in grey. In orange, events thought to be driven by increased *Ne*, in blue, those thought to be driven by reduced *Ne*.

**Figure 4.**
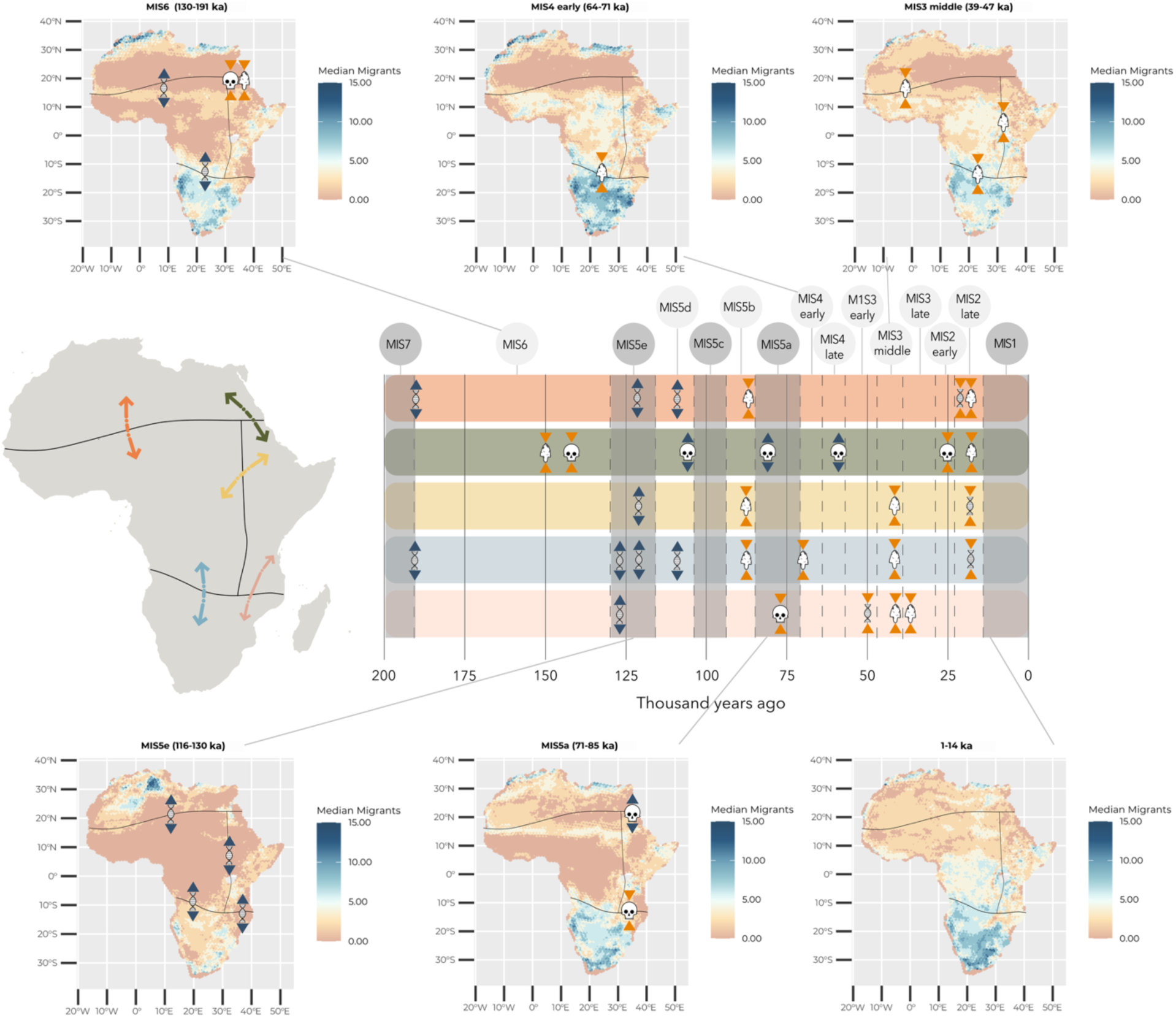
Migration over time and key events across regions. Maps are median effective migrants incoming to each deme from adjacent ones at each MIS stage for the Pan-African start scenario. Timeline indicates key genetic (DNA symbol), archaeological (stone tool symbol) and morphological (skull symbol) occurring across African regions (see Table S7), with interglacial periods shaded in grey. In blue, events thought to represent divergences between regions, in blue, those thought to represent convergences across regions.

Our model predicts an increase in fragmentation in the Eastern African range during MIS 5e and 5d (130-104 ka; Figs. 3 and 4), coinciding with a reduction in the corridor connecting this region to Southern Africa. This likely modulated the progressive divergence of the lineages leading to West and East Africans, a process estimated to have started ca. 130-120 ka^20,21^. This fragmentation would also explain the appearance of an unusual fossil in Lake Eyasi, Tanzania (Eyasi I, 130-80 ka)^82^ showing several archaic characteristics, despite the presence of anatomically modern *H. sapiens* much earlier in other parts of Eastern Africa^48,83^.

Although the Maghreb is thought to have been isolated from the rest of Africa for most of our evolutionary history^67,84^ (but see Text S1 for a discussion on how climate models might slightly underestimate rainfall in this area), fossils have been found during MIS 6, such as those from Rabat-Kébibat (137±7 ka) exhibiting archaic features similarly observed elsewhere in the Kabwe (299±25 ka) or Herto (157±3 ka) specimens^31,46,85–87^. The appearance of Aterian industries at Ifri n’Ammar (Morocco) at around 140 ka has also been hypothesised to stem from an existing pool of knowledge deriving from sub-Saharan populations that moved northwards during this period of increased connectivity^88–90^. This time also sees the oldest use of ornamental shell-beads in the region^91^. Our model contextualises these findings by showing that, despite the isolation of the Maghreb from sub-Saharan Africa as observed from the time-averaged plots (Fig 4), a more detailed reconstruction of changes through time reveals three main windows of increased connectivity and population sizes in the Maghreb at ca. 185 ka, 160 ka and 137 ka (Fig S27).

In our simulations, MIS 5c and MIS 5b underwent a progressive increase in estimated effective population sizes following a drop during MIS 5d. However, this process was not synchronous and started in Northern Africa in MIS 5c while Central Africa was still decreasing in population, with the population increase only reaching a substantial level in Central Africa in MIS 5b (Fig 3 and 4). Although, as mentioned above ornamental shell-beads had been used in Northern Africa since MIS6, an increased prevalence of these is observed in Morocco and Algeria during the period between 100 and 95 ka^9^. Similarly, in Central Africa, whilst some elements of the MSA blended with Early Stone Age were present during MIS6^92^, approximately 88 ka MSA technology without ESA elements and elaborate bone tools is found at Katanda 9 in the DRC^93–95^. Our model also predicts an increase in effective population sizes in Southern Africa 95 ka (MIS 5b), peaking around 80 ka (Fig 3; Fig S20), and coinciding with a period of higher innovation rates in the area exemplified among others by the appearance of engraved ochres in Blombos Cave^96^, engraved ostrich egg shells from Diepkloof^97^ and an increase in the frequency of bone artefacts in both Central and Southern Africa^98,99^.

MIS 4 also witnesses some key events in the evolutionary history of *H. sapiens* (Fig 3 and 4). At 67 ka, the site of Panga Ya Saidi (Kenya) experienced the earliest onset of the LSA in Africa^100^, and Southern Africa witnessed the appearance of the Still Bay and Howiesons Port industries characterised by a substantive increase in technological complexity^101,102^ argued by some to have required an increase in human cognitive capacities^101^. Instead, we find that these events coincide with the first window (70-65 ka, MIS 4) of reconstructed widespread Pan-African connectivity since MIS 7 (200 ka). Interestingly, the higher interconnectivity and metapopulation sizes of *H. sapiens* across Africa from the middle of MIS 4 onwards can also partly be attributed to a shift in *H. sapiens* preference/tolerance for more seasonal environments with higher precipitation (Fig S28). This shift might have been facilitated by cultural innovations and also partly explain the ability of *H. sapiens* to expand into regions previously rarely populated^38^ such as the Sahara, and ultimately leading to the global expansion of *H. sapiens*^2^.

Whilst after this period *H. sapiens* exhibits comparatively higher connectivity levels across the whole African continent, the onset of MIS 3 (54-48 ka) led to a temporary dip in movement across Central Africa (Fig S13). This dip provides an explanation for the estimated split time of the lineage leading to today’s Western (e.g. Aka, Baka, Bakola) and the Eastern Central African hunter-gatherers (e.g. Mbuti)^13,14,22^ (but see^34^), as well as highlighting a period during which the gene flow between the Aka and the San in Southern Africa would have been interrupted^103^.

In our model, connectivity between Central Africa and the rest of the continent picked up again by the latter part of MIS3 (39-29ka), when the earliest evidence for a transition from MSA to LSA industries is observed in eastern and western Central Africa at Matupi Cave (DRC)^104^ and Okala (Gabon)^105^, as well as in Southern Africa^106,107^. The disappearance of Aterian MSA technologies in North Africa which characterised the region since ca. 140 ka^30^ also takes place These changes are consistent with a period of increased demographic growth and connectivity across the continent fuelling innovation and cultural exchange. Genetic studies have hypothesised that during the mid-MIS 3, Mota-related ancestry spread as far south as Zambia, and Southern-African-related ancestry as far north as Kenya^14,23^, in line with our reconstructed increase in migration levels.

Finally, during the Last Glacial Maximum (LGM; 26-19 ka) in late MIS 2 we observe another expansion in the niche of *H. sapiens* towards environments characterised by greater precipitation and temperature seasonality (Fig. S28), as well as lower temperatures^38^. This niche expansion increases even further during the Holocene. The ability to occupy novel niches allowed *H. sapiens* to maintain high levels of connectivity and population sizes during most of the LGM and the Holocene across the continent, underlying both the transition towards anatomical modernity and the ubiquity of LSA technologies in Africa. Of note, however, is an increase in fragmentation in Western Central Africa during the Holocene, which supports the diversification of the lineages leading to the different Western Central African hunter-gatherer populations (ca. 12 ka; Fig S13)^108,109^.

By integrating paleoenvironmental, archaeological, morphological, paleogenetic, and genetic data within a single analytical framework, we built a Pan-African metapopulation model that reconstructs the demographic and evolutionary history of our species. Similarly to other studies^26,27,49^, our work indicates the presence of a deep African metapopulation established prior to 300-250 ka. Regardless of whether our models assume an initial expansion of *H. sapiens* within Africa, the demographic and genetic patterns across different scenarios converge on a single, coherent dynamic for the period after 200 ka, which we validated with independent anthropological, archaeological, morphological and genetic data. The demographic and connectivity changes inferred by this Pan-African metapopulation model provide a unified explanation for many of the key moments in the evolution of our species in Africa over the last 200,000 years.

Only after the establishment of structured populations in key regions of the continent would derived morphological, cultural and genetic traits observed in today’s members of our species have evolved (Table S7). Our results are consistent with findings that despite deep genetic divergences, there is evidence for intermittent episodes of gene-flow between all hunter-gatherer lineages^26,103^. Such episodes would have facilitated the emergence of adaptive cultural and phenotypic variants for local environments during periods of isolation on the one hand, and the exchange of such variants during periods of connectivity on the other^2,10,32,110^. Crucially, our model highlights the vast diversity of environments in which the members of our species thrived throughout our evolutionary history, and thus, the adaptive potential of the human foraging niche^111^.

The maintenance of viable population sizes and interconnectedness in most African regions following their initial settlement by early hunter-gatherers explains the evidence for low levels of historical inbreeding and high genetic diversity indicated by very few and short runs homozygosity observed among contemporary and ancient African hunter-gatherers^112^. It also aligns with archaeological^32,113^, ethnographic^110,114^ and paleoenvironmental evidence^56,115^ for the role of hunter-gatherer mobility and interconnectivity in human evolution, consistent with the idea that these populations are culturally adapted to promote and place value on these behaviours^116^. An exciting avenue for future research should investigate the role of culture in mediating the relationship between environmental and population dynamics. For instance, archaeological studies suggest that the niche expansion of *H. sapiens* at the onset of MIS 2 in East Africa was driven by cultural adaptations such as food storage, strategies that would have facilitated survival in increasingly seasonal environments during the LGM^117,118^.

In conclusion, our climate-informed reconstruction of the demographic history of *H. sapiens* provides quantitative evidence for a deeply structured Pan-African metapopulation undergoing repeated cycles of isolation and connectivity over at least the last 200,000 years. While we cannot entirely exclude the possibility of admixture with “ghost” archaic or “ghost” modern populations—especially prior to 300 ka^15,49^ we found that such admixture is not required to explain either the genetic or the morphological diversity observed across Africa over the last 200 ka^17,21,23,119,120^. Therefore, a Pan-African metapopulation model is sufficient to quantitatively account for the vast diversity observed among members of our species both in the past, as well as today.

The lack of consensus between current models relying purely on genetic data (and thus on a simplistic story of pulses of migration, population expansions or collapses at broad scales) might partly stem from the exclusion of population structure in hominin evolutionary models, which can lead to spurious inference of “ghost” admixture events^26,121^. Newly available data will hopefully open the way for an exciting avenue of research exploring the early establishment of a Pan-African metapopulation in the Middle Pleistocene^27,49^, and determine its relationship with other hominins inhabiting Africa and beyond^122,123^.

Moreover, whilst our model highlights the role of climatically driven changes in population dynamics in shaping continental-level archaeological transitions, future work combining our approach with models of the climatic drivers of particular cultural traditions^124^ may be instrumental for determining the selection pressures shaping the cultural evolution of our species.

## Materials and Methods

### Genetic data

Whole genomes from modern and ancient hunter-gatherers in Africa were obtained from publicly available databases.

For modern samples, we selected all hunter-gatherers available from the SGDP^125^, and HGDP^22^. We also selected genomes from populations that only recently abandoned a foraging lifestyle but are well known to have foraging ancestors, such as the Khomani San and Ju’hoansi from Namibia and South Africa or the Ogiek and Sandawe from Kenya and Tanzania. In addition, we also obtained genomic samples from agriculturalist populations from Eastern and Western Africa as well as from European individuals for the purpose of the genomic masking procedure we employed (see below). All the genomes have ∼30X as mean coverage.

For ancient individuals, we selected all available shotgun samples with a genomic coverage of ca. 1X and above (see Dataset S1 for initial genomes). We selected only those samples with a clear archaeological context associated with a foraging lifestyle based on the lack of presence of cultivated crops, metal or ceramic; and that subsequent genomic analyses confirmed their relatedness to contemporary and ancient foragers.

In total, and excluding Europeans and non-hunter-gatherer Africans, our sample comprised N=71 hunter-gatherer individuals belonging to N=26 populations, of which N=15 were ancient (Fig 1A; Fig S4).

### Processing of genetic data

For modern and high coverage ancient genomes (genomic coverage >10X), we downloaded all data in the form of FASTQ files, and followed the pipeline detailed in Maisano Delser et al.^8,126^ to call genotypes. The 10X threshold was chosen after a series of tests downsampling >30X genomes to determine whether calling genotypes pairwise pi results equivalent to those of the non-downsampled genomes (Fig S29).

The same tests also illustrated that genomes with an average genomic coverage of around 2X produced accurate pairwise pi results when, instead of genotyping diploid genomes, we took the consensus sequence of data to generate a pseudo-haploid genome. This enables us to avoid biases introduced by increased genotyping error of low-coverage data. To do so, we used ANGSD *-doCounts* v0.9^127^ to extract the frequency of mapped bases at each position of the mapped reads data filtering against low mapping (-minMapQ 20) and base calling (-minQ 20) after performing several checks to determine the average depth of coverage per position at different base qualities, and then used *consensify_c* v2.0^128^ to generate pseudo-haploid genomes. Compared to common pseudo-haplodifying strategies which normally sample a base from reads randomly, *consensify_c* allows for random sampling and majority rule calling. Here, we sampled 3 bases for each position in the mapped data, and called a consensus base when at least 2 of those agreed, filtering for positions with a minimum depth of 3. This strategy reduces the possibility of getting erroneous genotypes for low coverage data.

### Genomic masking of agriculturalist and European-related ancestries

Given our interest in reconstructing the evolutionary history of African hunter-gatherer genetic lineages, we had to remove genomic components that had been acquired through historical admixture with non-hunter-gatherer groups from the genomes of African hunter-gatherers^20,33,129^. To do so, we merged all modern genomes, removed invariant sites (using *bcftools*), and phased them using BEAGLE 3.0^130^ (and the same reference genome) with default settings. We then used PLINK v1.9^131^ to filter to maf=0.15 (given our sample size of N=84 genomes) and to perform LD pruning with a sliding window approach. Specifically, we evaluated SNPs in windows of 200 variants, shifted the window by 25 SNPs at each step, and removed one of any pair of SNPs with a squared correlation coefficient greater than 0.4. We then ran ADMIXTURE^132^ to estimate for every individual the proportions of the genome originating from K ancestral populations, K being specified a priori. The programme was run at K values from 3-10 with CV error estimation, default values for fold iteration (v=5) and a random seed for each K value. Therefore, for each value of K, we ran as many iterations as required for the log-likelihood to increase by less than ε=10^-4^ between iterations. We then inspected results and saw that at K=5, distinct clusters corresponding to European-related ancestries, African agriculturalist ancestries, Southern African hunter-gatherer ancestries, Eastern-Central African hunter-gatherer ancestries and Western-Central African hunter-gatherer ancestries appeared (Fig S24).

We used *Gnomix*^133^ on our dataset to perform semi-supervised local ancestry inference with references from K=4 ancestry clusters, corresponding to Central African hunter-gatherers (eastern and western), Southern African hunter-gatherers, European, and African agriculturalist populations using individuals that showed over 99% of ancestry from each of the respective groups. To try to balance reference panels, we kept the number of individuals per reference panel between 4 and 10 individuals. As recommended by the developers of the software, we used 15% of the data for training set 1, 80% for training set 2 and 5% for validation, a window size of 0.2cM, and an *r_admixed* parameter of 18 in *Gnomix*. Across chromosomes, the average classification accuracy of haplotypes to their correct ancestry was 95% (range=79-99%). This is much higher than what is normally achieved by other local ancestry inference methods^133^.

Before masking away chromosomal segments associated with African agriculturalist and European ancestries, we reintroduced the invariant sites into the re-phased *vcf* files obtained from *Gnomix* using *bcftools.* Invariant sites were assigned to the ancestry of origin associated with the 0.2 cM window where they belonged. Last, we removed African agriculturalist-associated chromosomal segments and European-associated segments^110,134^, in a procedure known as genomic masking. This means that positions located in genomic segments classified by *Gnomix* as belonging to those ancestries were masked (i.e. set to missing) for downstream analyses. We refer to the remaining unmasked chromosomal segments as hunter-gatherer chromosomal segments, and we refer to analyses that use only these segments as hunter-gatherer ancestry-specific analyses.

### Calculation of between-population pairwise pi values

After all modern and ancient genotypes had been called, we merged the low coverage ancient, high coverage ancient, modern African hunter-gatherer genotypes (with masked segments) and African agriculturalist genotypes in a single *vcf* file using custom-made scripts.

We selected 9178 loci of 10kb in length along the genome to represent neutral genetic variation^8,126^. We first applied filters to exclude regions that may be under selection or that could be problematic in terms of assembly quality: more specifically, we filtered out coding regions, conserved elements, recombination hotspots (regions with recombination rates >10cM/Mb), repetitive regions, and regions with poor mapping or sequencing quality. The filters are the same as used in Kuhlwilm et al.^135^ and Maisano-Delser et al.^126^ and were provided by Ilan Gronau. Contiguous intervals of 10kb were chosen on these remaining sites using a sliding window approach. Windows were retained if 7,500 sites or more were present in a panel of 7 modern samples distributed worldwide^126^, and a subset of these were selected with a minimum inter-locus distance of 50kbp. This minimum 50kbp distance between loci was chosen so that the chance of recombination was sufficiently high that loci could be treated as unlinked. Finally, 109 regions were discarded after filtering for coverage and CpG sites leaving 9069 windows.

We then determined the minimum number of windows, and the minimum number of overlapping (non-missing) positions that pairs of samples needed to have in order to allow us to calculate reliable whole-genome pairwise **π** (**π**_WG_) nucleotide differences by downsampling our reference genomes (unmasked) to a varying number of windows of a varying number of base pairs and calculating pairwise **π** values between them. We found that 2000 windows of 2000 base pairs each resulted in pairwise **π** values within 0.00006 of one another (Fig S30). We therefore only calculated **π**_WG_ between pairs of samples that shared over ≥2000 windows, each with ≥2000 overlapping base pairs. This forced us to exclude samples from contemporary Eastern African hunter-gatherers such as the Hadza, Sandawe and Ogiek as after masking away fragments associated with African agriculturalist and European ancestries, they did not share sufficient windows with the specified minimum number of positions with any of the remaining samples in our dataset. We also had to exclude three ancient samples from South Africa due to the same reason (BallitoBayB_2100BP, South_Africa_1200BP and South_Africa_2200BP).

For each window, we then calculated pairwise nucleotide differences between each pair of haplotypes (**π**_window_). For diploid samples compared with diploid samples, we took the average of 4 possible comparisons between haploids as the **π**_WG_ between the two samples for that window^126^. For haploid samples (those called with *consensify_c*), we individually considered all overlapping sites. The mean of all **π**_window_ across all windows was then used as empirical **π**_WG_ between samples.

**π**_window_ comparisons involving ancient genomes that were not Uracil-DNA Glycosylase (UDG) treated would have not been reliable because of the inflation of transitions in ancient genomes due to DNA damage and degradation. To circumvent this problem, we calculated the ratio of transitions (ts) to transversions (tv) across all samples and found as expected that samples from Mota_4500BP, BallitoBayA_1900, Morocco_7500, Matjes_River_5500, Matjes_River_3800, Matjes_River_7700, Cape_StFrancis_2200 and Springbokvlakte_1400 had unusual ts/tv ratios, which is expected given that they were not or only partly UDG treated. The average ts/tv ratio was calculated across all other samples and found to be 1.57. Note that, as expected, this ratio is significantly lower than the ts/tv ratio found across non-African populations given the reduced number of transitions occurring among Africans^8,136,137^. Then, estimates for the **π**_window_ between the three non-UDG/partly-UDG treated samples (**π**_window_from_tv_) and all other samples was calculated using exclusively transversions and multiplying the result by 1.57. We also assessed the correlation between **π**_WG_ and **π**_WG_ calculated only from transversions for modern and UDG treated samples to confirm the accuracy of this method (Fig S31).

Last, between population **π** (**π**_POP_) were obtained by averaging **π**_WG_ between all comparisons for each pair of populations.

### Dataset of hunter-gatherer archaeological sites

We expanded the dataset employed by Hallett et al.^38^ (N=479 sites) to include additional sites from Central Africa (included in^95^) as well as sites dated from the Holocene (14 ka until the present). As in Hallett et al.^38^ (which derives from a curated version of the ROCEEH Out of Africa Database (ROAD; last accessed 23/11/2021)^138^, we only included sites with 1) published coordinates, 2) radiometric dates, 3) an age error range less than or equal to 20,000 years. In addition, since after ca. 8 ka populations across the African continent start developing farming, yet we only wanted to model the time-dependent niche of hunter-gatherer populations, we excluded sites that presented evidence of cultivation, pottery, or metal use following the criteria detailed in Padilla-Iglesias et al.^56,95^. In doing so, we aimed to retain sites that were only occupied by hunter-gatherers. This resulted in a dataset of N=1242 African hunter-gatherer archaeological sites ranging from 120,000 cal BP to 495 cal BP (Dataset S2; Fig S1).

For the reasons outlined in Hallett et al.^38^, for all uncalibrated 14C ages entered in our database, a basic IntCal20^139^ calibration for northern hemisphere archaeological localities and a basic SHCal20 calibration for southern hemisphere archaeological localities was applied using the *rcarbon* R package^140^. The calibrated 14C ages were then used to determine mean age, and age error ranges were calculated to the level of 1σ. If an archaeological layer was dated with 14C and additional methods, our calibrated 14C mean age and 1σ error were then combined with the mean age and error range of the other methods to find the mean age used in our models.

### Ecological niche modelling of Homo sapiens and reconstruction of suitability landscapes

Our hierarchical modelling framework consists of model components that operate at different spatial and temporal scales (Table S8; Text S1). It is therefore necessary to transform and refine the environmental information entering our framework at the top (i.e., climate model simulations and their extension for 300 ka) into information suitable for the model component at each respective level in this hierarchy (e.g., ENM and CISGeM; see below). Although each data transformation likely introduces uncertainty into the data, the overall dimension of information being passed on from one level to the next is actually being reduced.

We modified the time-dependent ENM employed by Hallett et al.^38^ to quantify the extent of the suitable range of *H. sapiens* throughout the past 300,000 years. To do this, we used the R package *pastclim*^141^ to access the paleoclimatic simulations from Krapp et al^142^. These simulations are based on an emulator of the HadCM3 climate model, and includes 17 BIOCLIM variables (BIO 2 and BIO 3 are not available as they are based on daily rather than monthly summaries) and two vegetation variables (net primary productivity, NPP, and leaf area index, LAI). For the ecological modelling we used the same variables from Hallett et al.^38^: leaf area index (LAI), temperature annual range (BIO 7), mean temperature of the wettest quarter (BIO 8), mean temperature of warmest quarter (BIO 10), and precipitation of wettest quarter (BIO 16). In the original publication they were chosen as follows: when comparing distribution across the variable space in sites occupied by *H. sapiens* versus the whole of Africa in the period of interest (randomly drawing 10,000 points from each time slice), they showed the highest divergence. This can be interpreted as a specific ecological choice, which supports them to be the most informative to reconstruct habitat suitability for humans in this context^38^.

To take into account chronological uncertainty, we resampled each date 100 times from the truncated normal distribution identified by the mean and the plus/minus as 2-sigma^143^. Given the focus of the model by Hallett et al.^38^ on evaluating the possibility of a change in the niche occupied by *H. sapiens* over time, these authors had performed a thinning procedure to even out the number of presences in older and younger time periods. However, since for the present model we want to obtain the most accurate reconstruction we can, instead of reducing the number of occurrences in the same way, we only performed a spatial thinning to only keep one presence for every 200 km radius to prevent spatial autocorrelation of results^144^. To adequately represent the existing climatic space (i.e. background) in our ENMs, each of these resulting datasets was coupled with a random sampling, for each observation, of 200 random locations matched by time. This resulted in n=100 datasets (“repeats”) of differently sampled and dated presences and differently sampled background points which we used to repeat our analyses to account for the stochasticity of the sampling performed. We then followed Hallett et al.^38^ and used GAMs (*mgcv* package in R^145^) to fit two possible models: a “time-varying-niche” model that included interactions of each environmental variable with time (fitted as tensor products), and a “fixed-niche” model which included the environmental variables as covariates but lacking interactions with time.

Following Hallett et al.^38^, we performed standard checks on the residuals of the models using the DHARMa package in R^146^, including Kolmogorov–Smirnov (KS) tests for correct distribution, dispersion, and outliers.

For each repeat, we verified, through the use of Akaike Information Criterion (AIC) that the time-varying niche models were best supported (Fig S32). We then evaluated the fit of the changing niche models with the Boyce Continuous Index^147^ (BCI), a measure of model performance (calculated as a Pearson correlation coefficient) specifically designed to be used with presence-only data, setting an acceptance threshold > 0.7^148^ (Fig S33).

We wanted to model hunter-gatherer evolutionary dynamics in Africa from 300 ka until the present, but, because of data availability, our ENM could only be built for the period spanning between 120-1 ka. Therefore, to extend our model from 120 ka until 300 ka, we had to project model predictions to this older time frame. Projection is a common procedure in ENMs^149^, but as our model is built with a time-dependent relationship between environmental variables and habitat suitability we had to take such temporal changes into account. We tested projecting to 300-121 ka using the relationship between environment and time obtained for the time slices of 105, 110 and 120 ka. We observed that projections using the 120 and 105 ka relationships between environment and time were more consistent with one another than the one using the relationship at 110 ka. Therefore, we projected the 105 ka model from each repeat to create the suitability maps over the whole of Africa from 300-121 ka. We used the same procedure to project the time-dependent suitability maps for 1 ka to the present.

To achieve more robust results, it is common practice in ENM to summarise the predictions obtained from different repeats (or runs) as so-called “ensembles”^150^. In our case, we calculated the median ensemble and evaluated the results following the same procedure presented in Hallet et al.^38^ (Table S9).

Although we used normalised ENM predicted probabilities as inputs for CISGeM, to visualise the output of the ENM, we followed Hallett et al.^38^. We transformed the predicted probabilities of occurrence of the ensemble into binary (presence/absence) by using three thresholds based on the number of human presences (considering all resampled dates) encompassed. The basic idea is that some of the occurrences or resampled dates could be climatic outliers^38^. For this reason, predicting as suitable for humans the whole range of climate where sites have been observed (“total area”, encompassing 99% of occurrences) may result in an overestimation of the spatial distribution. So, we identified as more likely suitable, but still potentially overestimated (“peripheral”) or minimum predicted area encompassing 95% of our presences. Finally, we calculated as most likely suitable (“core”) the even smaller area that encompasses 90% of our presences. This was done using a modified version of the function *ecospat.mpa()* from the *ecospat* R package^151^. To better visualise how the reconstructed potential distribution changed through time, we then merged rasters from each period considered by calculating their mean.

### Comparison with previously published ecological niche models

We compared the results of our ENMs with the ones obtained by Hallett et al.^38^ calculating the overlap between the respective results using two common metrics used for this aim. We calculated Schoener’s D^152^ and a modified Hellinger metric^153,154^ to calculate the overlap between the *H. sapiens* niche estimated with our model (without resampling archaeological sites), and the model from Hallett et al.^38^ were presences at each time period were resampled to ensure an even number across time periods and built with paleoclimate reconstructions based on the HadCM3 model without an emulator. We used as human niche the binary predictions encompassing both the core and peripheral area of H. sapiens (90 and 95% of presences). We obtained a Schoener’s D value of 0.83 and 0.81, respectively, and Hellinger metric values of 0.95 and 0.95 respectively, which indicate very high niche equivalency^153,154^.

We assessed the robusticity of our ENM models to different climate models by comparing our ENM with the second model presented in Hallett et al.^38^ and built with the climatic dataset presented in Yun et al.^155^, based on the CESM climate model. The similarity between the two results calculated with the median Sorensen similarity Index across time periods^156^ was 0.85.

### Climate Informed Spatial Genetic Model

Reconstructed suitability maps for the last 300,000 years were used as inputs for CISGeM, a custom-made framework that stands for Climate Informed Spatial Genetic Models. This framework allows the modelling of the genetics of multiple populations within a spatially explicit reconstruction of the world, where the suitability of each deme changes through time according to scores from the ENM. Then, using a DL approach based on CNN, we fitted basic demographic parameters such as population growth rate and migration, as well as the link between ENM predictions and local population sizes. We also fitted costs to migration due to altitudinal changes. We calculated five summary statistics from the matrices of pairwise genetic differentiation (**π**) between all possible pairs of populations (see Materials and Methods) that were fed to the model, thus reconstructing in full the genetic population structure observed at the corresponding time where the genome was sampled.

CISGeM operates by simulating the expansion dynamics of a population from a position of a given area in a map (Fig S2)^8,36^. Since we were simulating the dynamics of early Anatomically Modern Humans (AMHs) in Africa, and the place of origin of *H. sapiens* within the continent is far from clear^9,10,46,48^, we fitted four different start scenarios in CISGeM: One with a starting point in the Kibish Formation in Eastern Africa (Coordinates: 35.96, 4.80), another one with a starting point in the Jebel Irhoud site in Northern Africa (Coordinates: −8.86, 31.89), another one with a starting point in the Florisbad in Southern Africa (Coordinates: 20.07; −28.77) and another one where the simulation started by seeding all connected viable demes across the continent (a Pan-African start). The same procedure was used to run all four scenarios.

### CISGeM demography

CISGeM’s demographic module consists of a spatial model that simulates long-term and global growth and migration dynamics of AMHs. These processes depend on eight parameters (Table S1), which we later estimate statistically based on empirical genetic data.

The model operates on an equal-area hexagonal grid of 2,627 cells that represent the area of the African continent (the distance between the centres of two hexagonal cells is 120.6 ±7.6 km; the variation is due to the earth having a spheroid rather than a perfect spherical shape and thus the grid not being perfectly regular). Each time step represents 25 years, appropriate as a conservative estimation of the generation time in AMHs (^157^; see^158–160^ for alternative estimates). The specific settings used in our ENMs (especially the ratio between presences and pseudoabsences, which is 1/200) led to our suitability values to be extremely low (which is expected in such cases). To use them as continuous values within CISGeM’s hexagonal grid, we simply multiplied them by 100,000 to transform them into integers which are more easily handled by CISGeM. We then used the *-remaplaf* function from the Climate Data Operator (CDO) software (v3.2.0)^161^ to turn the square grid into a hexagonal grid.

Each time step of a simulation begins with the computation of the carrying capacity of each grid cell, i.e. the maximum number of individuals theoretically able to live in the cell for the environmental conditions at the given point in time. The carrying capacity (*K*(*x*, *t*)) (in effective individuals, *N_e_*) in a grid cell *x* at a time *t* was modelled as:

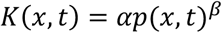

Where *p*(*x*, *t*) denotes the normalised ENM probability of *H. sapiens* inhabiting cell *x* at time *t* (see ENM section). *α* and *β* are two parameters: allometric scaling factor, and allometric scaling exponent, respectively, that determine the relationship between normalised ENM probabilities and carrying capacity. The framework does not propagate raw climate fields. It transforms the climate into representations aligned with the spatial and temporal sensitivities of ecological and demographic processes. We confirmed that this relationship is biologically plausible by verifying that, when fitted to the genetic pairwise **π**, it provides a good match to census population sizes for modern hunter-gatherer groups^55^ (see below). However, note that the parameters were fitted to the genetic data (see below); the census population sizes of modern hunter gatherers were only used to test that the shape of the relationship was realistic.

The estimated carrying capacities are used to simulate spatial population dynamics as follows. We begin a simulation by initialising a population in a group of adjacent cells (determined by the initial deme radius parameter) **x*_0_* at a point in time *t_0_* with *K*(*x_0_*, *t_0_*) individuals. Depending on the scenario we were running, *x_0_* was located in Eastern Africa (cell closest to 35.97, 4.80 degrees), Northern Africa (cell closest to −8.86, 31.90 degrees), Southern Africa (26.06, −28.77 degrees) or the whole continent. At each subsequent time step between *t_0_* and the present, CISGeM simulates two processes: the local growth of populations within grid cells, and the spatial migration of individuals across cells. Similar to previous work^162,163^, we used the logistic function to model local population growth in humans, estimating the net number of individuals by which the population of size *N*(*x*, *t*) in the cells a *x* at time *t* increases within the time step as

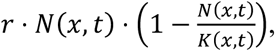

where *r* denotes the intrinsic growth rate. Thus, growth is approximately exponential at low population sizes, before decelerating, and eventually levelling off at the local carrying capacity.

Across-cell migration is modelled as two separate processes, representing a non-directed, spatially uniform movement into all neighbouring grid cells on the one hand, and a directed movement along a resource availability gradient. For both of them, movement between two grid cells is reduced when it involves crossing mountains. Under the first movement type, the number of individuals migrating from a cell *x_1_* into a neighbouring cell *x_2_* is estimated as

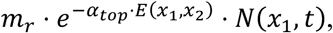

where *α*_top_, *t*_0_, *m_r_*, are parameters, and where *E*(*x_1_*, *x_2_*) is a measure of the altitude that needs to be covered between cells *x_1_* and *x_2_*, which we defined as follows. We used a very-high-resolution (1 arc-minute) global elevation and bathymetry map (ETOPO1) and determined, for each pair of neighbouring cells *x_1_* and *x_2_*, the altitude profile along the straight line between the geographic centres of the two cells. We then defined *E*(*x_1_*, *x_2_*) as the sum of the absolute values of all altitude changes along the line. This assumes that descends have the same effect in terms of reducing movement rates as ascends; in particular we have *E*(*x_1_*, *x_2_*)=*E*(*x_2_*, *x_1_*). This mechanism in Eq. (1) is equivalent to a spatially uniform diffusion process, which has previously been used to model random movement in AMHs^162,163^. Under the second movement type, an additional number of individuals moving from a grid cell *x*_*_to a neighbouring cell *x*_+_is estimated as

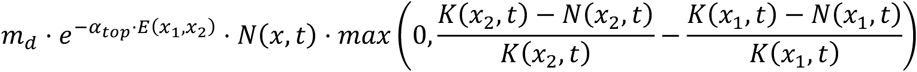

The number 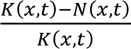 represents the relative availability of unused resources in the cell *x* at time *t*, equalling 1 if all natural resources in *x* are potentially available for humans (*N*(*x*, *t*) = 0), and 0 if all resources are used (*N*(*x*, *t*) = *K*(*x*, *t*)). Thus, individuals migrate in the direction of increasing relative resource availability, and the number of migrants is proportional to the steepness of the gradient. The distinction between directed and non-directed movement allows us to examine to which extent migration patterns can be explained by random motion alone or requires us to account for more complex responses to available resources (it is not uncommon in population genetics models to distinguish between colonisation and migration proportions, which usually differ when fitted to data). The coefficient *m*_3_ is a parameter.

It is in principle possible that the number of individuals present in a cell after all migrations are accounted for (i.e. the sum of local non-migrating individuals, minus outgoing migrants, plus incoming migrants from neighbouring cells) exceeds the local carrying capacity. In this case, incoming migrants are rescaled proportionally so that the final number of individuals in the cell is equal to the local carrying capacity. In other words, some incoming migrants perish before establishing themselves in the destination cell, and these unsuccessful migrants are not included in the model’s output of migration fluxes between grid cells. In contrast, non-migrating local residents remain unaffected in this step. They are assumed to benefit from a residential advantage^164^, and capable of outcompeting incoming migrants.

CISGeM’s demographic module outputs the number of individuals in each grid cell, and the number of migrants between neighbouring grid cells, across all time steps of a simulation. These quantities are then used to reconstruct genetic lineages.

### CISGeM predicted genetics

Once a demography has been generated, gene trees are then simulated (the genetic code borrows heavily from *msprime*^165^). This process depends on the population dynamics recorded during the demography stage and assumes local random mating within cells according to the Wright-Fisher dynamic. From the present, ancestral lineages of sampled individuals are traced back through generations, recording which cell each lineage belongs to. At every generation, the lineages are randomly assigned to a gamete from the individuals within its present cell. If the assigned individual is a migrant or coloniser, the lineage is moved to the cell of origin for that individual before ‘reproduction’. Common ancestor events happen when two lineages are assigned to the same parental gamete and they are then merged into a common ancestor lineage. This process is repeated until all the lineages have merged. If multiple lineages are still present at the time when the demography was initialised (either in a single deme, or in multiple demes), the remaining lineages enter a single ancestral population (with fixed population size *K_0_*), and a coalescent model is used to estimate the timing of additional common ancestors events to close the tree (see^166^ for an example of using hybrid models where the coalescent is used to close trees generated by an initial Wright-Fisher dynamics).

### DL-CNN architecture

We developed our DL-CNN architecture using Keras v3.10.0^167^ with a TensorFlow v2.19.0^168^ backend to automatically extract information from a set of five features (mean, median, maximum, standard deviation and median absolute deviation) calculated over 2000 simulated genomic windows for all possible 161 pairwise **π** among populations. The simulated data is then loaded into a CNN^39^ that begins with a projection layer with a 1D convolution with 16 filters and a kernel size of 1. This operation linearly combines the 5 features into a higher-dimensional representation without mixing the spatial information across different pairwise comparisons. This is followed by two blocks of 1D convolution layers (the first with 32 and the second with 64 filters) and a kernel size of 3 for spatial feature extraction, to capture local dependencies across the 161 genomic windows. All three convolutional layers were followed immediately by batch normalisation. Each of the last two convolutional blocks is followed by a max-pooling layer (pool size = 2 and 3, respectively) to reduce the dimensionality of the feature maps. The resulting feature maps are flattened (collapsed into a single linear vector) and fed into a fully connected (dense) layer containing 64 neurons with a dropout layer with a rate of 0.5 to prevent overfitting.

We used the nonlinear Swish activation function^169^ in all hidden convolution and dense layers. The network parameters were updated using the Adam optimiser^170^. The model was trained for a maximum of 128 epochs with a batch size of 32. To further prevent overfitting, we implemented an early stopping mechanism that monitored the validation loss with a patience of 50 consecutive epochs, and saved the epoch with lower validation loss. The loss function was different depending on whether the network was being trained for parameter estimation or scenario classification. The Mean Absolute Error loss function was used for parameter estimation, while a categorical cross-entropy loss function with label smoothing (alpha=0.3, to reduce overconfident predictions) was used for scenario classification.

We included both aleatoric and epistemic uncertainty to estimate the posterior distributions of the empirical data. For aleatoric uncertainty, we used 100 iterations of a bootstrap resampling strategy that consisted of sampling with replacement 2000 random genomic windows from the full 9069 windows in the empirical dataset. To account for epistemic uncertainty, for each of those replicates, we applied 100 Monte Carlo Dropout^171^ stochastic forward passes.

### Parameter estimation and model fitting

Parameter space was explored with a Monte Carlo sweep, in which demographic parameters were randomly sampled from flat prior ranges (Table S1). A fixed mutation rate of 1×10^-8^ was used.

We generated, for each scenario, the following number of simulations:

- Northern Africa: 56465; 31039 reached all samples
- Eastern Africa: 55352; 31663 reached all samples
- Pan-African: 66120; 37654 reached all samples for model comparison and 90079; from which 51330 reached all samples for parameter estimation.
- Southern Africa: 60029, 31113 reached all samples

Parameter estimation was performed with the DL-CNN approach described above. For each model, we trained the network using 50,000 simulations for the Pan-African scenario and 30,000 simulations for the other scenarios, which were split into a training and a validation set (with 80% and 20% of those simulations, respectively). We performed a power analysis by using 1,000 independent simulations (not seen during training) from each scenario as a test set. For each of the 1,000 simulations of the test set, we assessed the predictive power by comparing the true parameter values with the posterior predictions, calculating the coefficient of determination (R²). Results indicated good power (R² > 0.15) for all parameters in models with different starting demes, including the allometric scaling factor, allometric scaling exponent, intrinsic growth rate, undirected and directed expansion coefficients, altitude cost, and initial time (Table S2). For the Pan-African start scenario, the parameters directed expansion and intrinsic growth rate showed low power (R² < 0.05), while all other parameters had high power (R² > 0.7). We plotted the obtained posterior distribution of the parameters (obtained by including both epistemic and aleatoric uncertainty by using the Monte Carlo Dropout and bootstrap resampling strategies, respectively) when the empirical data was used for predictions (Fig S8-S11).

We then assessed whether the simulations were able to recreate the observed genetic diversity by comparing the ranges of the 5 summary statistics obtained for all 161 population pairs and verifying that the observed values for all of them fitted within the range simulated by the model. We also looked at the 161 average pairwise **π** singular values to further validate the goodness-of-fit between our simulations and the empirical data (Fig S5). In addition, to make sure our model captured all key aspects of the genetic patterns simultaneously, we used the first 2 principal components of each of the five summary statistics (explaining at least 65% of the variance), to verify that scenarios were deemed capable of reproducing genetic diversity if the observation was within the 2-dimensional cloud of predicted values on the 2 PC axes (Fig S6).

### Model comparison

To assess the relative fit of the scenarios with the different starting demes, we also used the DL-CNN approach described above and changed the output layer to perform classification rather than parameter regression. We trained the network with the same five simulated summary statistics from the simulations from each of the scenarios - using the summary statistics as predictor variables and the scenario labels as response variables. Then, we classified the observed 161 pairwise **π** comparisons among populations using the trained network, and obtained the probabilities (also using the Monte Carlo Dropout and bootstrap resampling strategies to account for uncertainty) of the observed data belonging to each of the scenarios. These probabilities were used to determine the relative likelihood of the observed data under each competing scenario.

### Demographic scenarios

We used the best 1000 simulations from each scenario based on the sum of Euclidean distances between observed and simulated mean summary statistics, to re-run CISGeM extracting demography output files every 5 generations (125 years). For each simulation, we extracted the values of effective population size, per cell, and the number of migrants between each pair of connected cells, per 5 generation period. We then obtained the median effective population size per cell, per 5 generation period, as well as the median number of migrants per triangle, per 5 generation period across the 1000 best simulations for each model.

Last, to determine demographic fluctuations over time, we used the median demography per 5 generation period to calculate the geometric mean in effective population size per grid cell over the entire duration of the simulation (Figure 2). To do this, we excluded cells that were uninhabited for over 50% of the duration of the simulation.

### Tests of population continuity in West Africa

Although there are no ancient nor modern genomes from Western African hunter-gatherers, the Yoruba, who live in Nigeria, have been hypothesised to descend from a “Basal Western African” lineage that separated from the lineage to which East African hunter-gatherers trace their ancestry, after Central, Southern and Eastern African lineages separated from one another^17^. Yet, other studies hypothesise that the ancestors of the Yoruba only recently migrated to Western Africa, from Eastern Africa^23^. Hence, whether the Yoruba are directly descendants of a Basal West African population, and the timing of the colonisation of Western Africa by this population remains unknown. To address this question, we sampled 30 random points in a regular grid covering the entire African continent, and placed a virtual population in each of the 30 points with the genetic profile of the Yoruba. We then re-ran CISGeM using the best fitting 1000 simulations from each start scenario and calculated, for each of the 30 samples, the mean difference between the simulated pairwise **π** between that population and every other “real” population, to determine which samples were located in the geographic position where our model would predict the Yoruba to be situated if they could be modelled as a hunter-gatherer population.

### Tests of relationship between contemporary hunter-gatherer population density and simulated effective population density

Padilla-Iglesias et al.^56^ provided evidence that delineating the niche of contemporary hunter-gatherers was informative for making predictions about patterns of past habitation. Hence, to verify the accuracy of our models, we used data on the geographic location of the N=749 camps from Central African Hunter-Gatherer populations compiled by Padilla-Iglesias et al.^56^ as well as a newly compiled dataset of N=562 additional camps from other populations of Eastern and Southern African hunter-gatherers from the literature (Dataset S4). Since CISGeM produces an effective population size estimate for each grid cell, we then used our hunter-gatherer camp dataset to obtain the indices of the N=519 cells that had camps in them, and compared the median predicted effective population size for the present across the best 1000 runs from each model (see above) of those cells to N=519 random cells from the continent. As an additional check, we also compared the mean BCI from the cells with camps from our ensemble ENM to an equal number of randomly sampled cells.

We additionally compared our median predicted effective population size for the present with estimates based on a modelled surface of census sizes derived from N=358 ethnographic societies^55^ using a correlation test.

### Test of relationship between simulated genetic distances and morphological distances based on cranial morphology

Given that our aim was to assess the ability of our model to reconstruct *H. sapiens’* evolution at timescales deeper than were spanned by our genomes, we compiled a dataset of *H. sapiens* of crania from the 200 ka until the present (Dataset S3). We chose to focus on this timescale as it represents the time frame for which the entire African continent was colonised by *H. sapiens,* according to our model. We then obtained all those cranial measurements for which variability was not determined by climatic conditions among contemporary populations (see^57,58^), and thus, thought to be primarily driven by drift. After sensitivity analyses, we excluded specimens for which there were less than 5 of such measurements available, as well as measurements that were available for less than 50% of the specimens. We imputed the remaining measurements for each cranium using the k-Nearest Neighbour method^172^, which has been shown to produce the most accurate estimates in biodistance analysis in paleoanthropology (and specifically in the imputation of cranial measurements when missing data=50%^173^). We used k=5, which has been shown to be appropriate for the imputation of these types of data^173,174^. This left us with a final dataset comprising 45 crania belonging to 14 populations (defined as being recovered at the same geographical location and assigned a similar age), with 11 measurements per crania. We then adjusted the measurements for size differences between the specimens using the logged version of the division of each variable by the geometric mean of all variables for that specimen (c.f.^175^). Given that the dating of one of the specimens (LH-18) has been contested - some situate it at 200-300 ka but others at 129 ka^176^, we re-did analyses excluding LH-18, but results were qualitatively similar.

We then performed a PCA with all specimens and calculated Euclidean distances (henceforth morphological distances) between the specimens using the first 7 principal components. We chose this number of components after checking using a scree plot that it was after the 7th component that a threshold in the cumulative variance explained by each subsequent PC was reached (explaining >80% of the variance in the data).

Using the 1000 best fitting runs from the Pan-African, Eastern, Southern and Northern Africa start scenarios, we placed samples at the geographic location of each of the crania in the dataset at its corresponding time period, and ran CISGeM. Then, we calculated the mean simulated pairwise **π** between each pair of crania across the 1000 runs for each model, and compared them to the observed morphological distances using Mantel tests in the R package *vegan*, with 10,000 permutations^177–179^.

We further calculated the great-circle geographic distance between cranial locations as well as cost-adjusted distances. Cost-path distances between sites were derived from the 1 km SRTM DEM using Tobler’s hiking function implemented in the *gdistance* package^180^. To evaluate the relationships among morphological, simulated genetic, and spatial distances, we tested for associations between morphological distance, geographic distance, and cost-adjusted distance.

In addition, we performed partial Mantel tests to assess the relationship between simulated genetic and morphological distances while controlling for (i) geographic distance and (ii) cost-path distance separately. This allowed us to verify that any observed correspondence between simulated pairwise **π** and morphological distances was not simply driven by isolation by distance.

## Data availability

All the genomic data used in this manuscript are deposited in public servers. See Dataset S1 for accession numbers of each of the samples. Archaeological data to build the ecological niche model are available in Dataset S2. Fossil cranial specimens used for morphological analyses are available in Dataset S3.

Contemporary hunter-gatherer camps used for model validation are available in Dataset S4 and Padilla-Iglesias et al.^56^. All paleoclimatic data used is available from the R package *pastclim* (https://github.com/EvolEcolGroup/pastclim). Data from Tallavaara et al.^55^ used to validate our demographic reconstructions was obtained from the authors of the original study.

## Code availability

All the code required to reproduce the analyses reported in this manuscript, as well as instructions on how to do so are available in the following GitHub repository: https://github.com/EvolEcolGroup/African_HunterGatherers; which in turn relies in the program *CISGeM* (https://github.com/EvolEcolGroup/cisgem) and the R package *rcisgem* (https://github.com/EvolEcolGroup/rcisgem). All three repositories will be made public upon acceptance of the manuscript.

## Acknowledgements

C.P-I. wishes to acknowledge Luisa Espinós Iglesias for help with the manuscript figures. C.P-I was funded by Fundación La Caixa and Forschungskredit Candoc (grants LCF/BQ/EU19/11710043 and FK-19-083) and Emmanuel College, Cambridge. M.L. and A.M. were funded by the Leverhulme Research Grant RPG-2020-317. M.F.P. was funded by the Talent Attraction Program of the Madrid Community, Spain (2024-T1/ECO31482). J.L.A.P. was funded by the Marie Skłodowska-Curie individual fellowship “RESOURCEFUL” (101028348). A.H. was funded by the Marie Skłodowska-Curie individual fellowship “FeverTime” (101063265). E.M.L.S. and M.C. were funded by the Lise Meitner Pan-African Evolution Research Group. P.M.D., M.L., M.C., and A.M. were supported by ERC Consolidator Grant “LocalAdaptation” (647797).

## Author contributions

Project conceptualisation: C.P.-I., A.M.; Project supervision: A.M., L.V., A.B.M.; Funding acquisition: A.M., E.M.L.S., A.B.M., C.P.-I.; Data acquisition: E.Y.H., C.P.-I., M.L., I.C., K.L., J.N.C., A.W.K., G.L., E.M.L.S., M.W.; Methodology: Z.X., M.L., J.L.A.P., M.C., M.K, A.H., P.M.-D., J.B.-P., A.G.I., A.M., C.P.-I.; Formal analysis: A.M., M.L., C.P.-I., M.F.P.; Writing (original draft): C.P.-I., A.M.; Writing (review & editing): all authors. All authors provided useful comments and guidance throughout the project.

